# A genetically engineered therapeutic lectin inhibits human influenza A virus infection and sustains robust virus-specific CD8 T cell expansion

**DOI:** 10.1101/2024.08.15.608041

**Authors:** Meng Yu, Ang Lin, Faezzah Baharom, Shuijie Li, Maureen Legendre, Evelyn Covés-Datson, Ebba Sohlberg, Susanne Schlisio, Karin Loré, David M. Markovitz, Anna Smed-Sörensen

## Abstract

Native banana lectin (BanLec) is antiviral but highly mitogenic, which limits its therapeutic value. In contrast, the genetically engineered H84T BanLec (H84T) is not mitogenic but remains effective against influenza A virus (IAV) infection in mouse models. However, the potency and effect of H84T on human immune cells and IAV-specific immune responses is undetermined. We found that H84T efficiently inhibited IAV replication in human dendritic cells (DCs) from blood and tonsils, which preserved DC viability and allowed acquisition and presentation of viral antigen. Consequently, H84T-treated DCs initiated effective expansion of IAV-specific CD8 T cells. Furthermore, H84T preserved the capacity of IAV-exposed DCs to present a second non-IAV antigen and induce robust CD8 T cell expansion. This supports H84T as a potent antiviral in humans as it effectively inhibits IAV infection without disrupting DC function, and preserves induction of antigen-specific adaptive immune responses against diverse antigens, which likely is clinically beneficial.

## INTRODUCTION

Influenza, caused by the influenza A virus (IAV), continues to cause significant mortality and morbidity worldwide. Seasonal influenza epidemics result in millions of infected individuals and cause 500,000 deaths annually. In addition, at irregular intervals the virus undergoes changes that can cause pandemics with high mortality ^1^. Virus-specific immune responses are critical to control and clear the infection, but if not properly controlled, may also contribute to immunopathogenesis and development of more severe disease ^2^. In addition, IAV has developed strategies to escape immunological control ^3–8^. Influenza vaccines contribute significantly to reduced morbidity and mortality. However, vaccine effectiveness is impacted by the ever changing nature of the virus and influenza vaccines are known to be less effective among the elderly ^9,10^. Thus, potent influenza-targeting antiviral agents would increase our ability to effectively prevent and/or treat influenza. Currently, there are only a few anti-influenza drugs in use clinically ^11,12^, and IAV strains resistant to these drugs are emerging worldwide ^13–15^. Therefore, identification and development of novel anti-IAV therapies would be desirable to expand the toolbox available to combat influenza.

Lectins are a group of proteins characterized by their ability to recognize and bind to specific carbohydrates without modifying them ^16^. Banana lectin (BanLec) in its native (wild type) form has broad antiviral effects against many enveloped viruses including IAV ^16^. BanLec restricts IAV replication *in vitro* ^17^ mainly via specific binding to high mannose bearing glycoproteins present at high densities on the surface of influenza virions ^18,19^. However, clinical applications of BanLec were long considered impossible due to the strong mitogenic properties of the lectin with potential systemic inflammatory side effects ^20,21^. A single amino acid substitution at position 84 of wild type BanLec (WT) from histidine to threonine (H84T), was shown to uncouple the mitogenicity from the antiviral activity of this lectin ^12,17^. By binding to hemagglutinin (HA) and inhibiting virus-endosome fusion, genetically engineered H84T restricts replication across IAV strains, including drug-resistant strains, in mammalian cells ^12^. *In vivo*, H84T protects mice from lethal IAV infection without causing mitogenic effects ^12^, indicating that H84T could potentially be applied as a broad-spectrum anti-influenza agent. However, to evaluate the clinical usefulness of H84T it is critical to determine the yet unstudied impact of the lectin on human immune cells and their function during viral infection. Dendritic cells (DCs) are innate immune cells that are well equipped to sense incoming viruses. DCs line the respiratory mucosa and are also recruited there in response to IAV infection ^22^. Importantly, DCs are antigen-presenting cells with the unique capacity to activate naïve T cells and are therefore critical for initiating the expansion and activation of antigen-specific T cells that are necessary to control and clear infection ^5,22^. We have previously shown that IAV infection impairs the capacity of DCs to present antigen to T cells, providing a partial explanation to why IAV infection is not only disease-driving on its own but also increases the risk of the secondary bacterial infections commonly seen in IAV patients ^5^. Here we set out to determine the effects of BanLec on human DCs during IAV infection, including their ability to present viral antigen to and activate T cells, to evaluate the future use of H84T as an antiviral treatment in humans.

## RESULTS

### WT BanLec, but not H84T, induces human T cell proliferation via dendritic cells in an HLA-DR- and CD86-dependent manner

We first confirmed that, as previously shown ^17^, H84T, unlike WT BanLec (WT), did not induce significant CD4 or CD8 T cell proliferation in human peripheral blood mononuclear cells (PBMCs) (**Figure 1A-B**) as compared to untreated PBMCs or PBMCs treated with D133G BanLec (D133G), a control lectin engineered to lose both antiviral and mitogenic capacity (**Figure 1A-B**). In contrast, in new experiments using only purified T cells, CD4 and CD8 T cells did not proliferate in response to BanLec (**Figure 1A-B**). Thus, we hypothesized that antigen-presenting cell (APC) - T cell interactions, present in PBMCs, may be important in mediating the mitogenic effect of BanLec. To test this hypothesis, we co-cultured autologous monocyte-derived dendritic cells (DCs) and purified T cells and found that WT induced significant CD4 and CD8 T cell proliferation compared to D133G or no BanLec (**Figure 1C-E**), restoring the pattern observed in PBMCs. Again, H84T did not induce T cell proliferation (**Figure 1C-E**), confirming lack of mitogenicity. Furthermore, blocking HLA-DR or CD86 in the DC-T cell co-cultures significantly decreased CD4 T cell proliferation induced by WT compared to the isotype control (**Figure 1F**), while only CD86 blocking significantly reduced WT-induced CD8 T cell proliferation (**Figure 1G**). Transfer of BanLec-stimulated DC supernatants did not induce T cell proliferation (**Figure 1H-I**), showing that the mitogenic effect of WT was dependent on DC-T cell contact. In summary, our data show that the mitogenic effect of WT is cell contact dependent in an HLA-DR- and CD86-dependent manner, while H84T displays no mitogenic effect on neither human PBMCs nor purified T cells.

**Figure 1.**
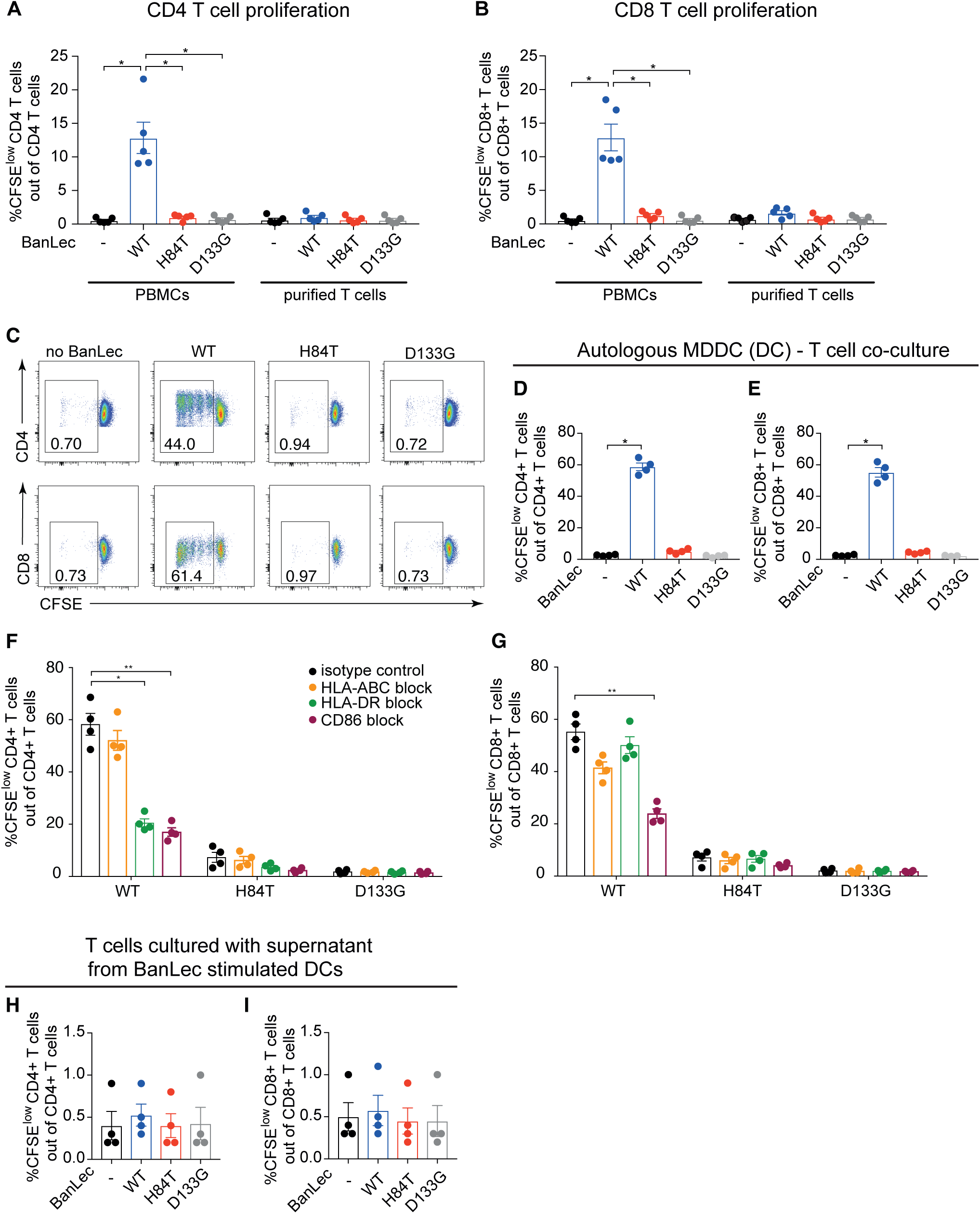
WT but not H84T, induces human T cell expansion by interacting with dendritic cells (DCs) in an HLA-DR and CD86-dependent manner. (**A-B**) CFSE labelled PBMCs or isolated T cells from the same donor were exposed to nothing (black) or to 2 μg/mL WT (blue), H84T (red) or D133G (grey) BanLec for 72 hours. T cell proliferation was detected by CFSE dilution using flow cytometry. Graphs show frequencies of live proliferating CFSE^low^ (**A**) CD4 T cells and (**B**) CD8 T cells from five individual donors with mean ± SEM. (**C-E**) DCs were exposed to nothing or 2 μg/mL WT, H84T or D133G for 1 hour, and then co-cultured with CFSE labelled autologous CD4 or CD8 T cells (1:10 ratio MDDC:T cells) for 5 days. T cell proliferation was detected by CFSE dilution using flow cytometry. (**C**) Dot plots show live CD4 or CD8 T cells and numbers indicate frequency of proliferating CFSE^low^ T cells. One representative donor of four is shown. (**D-E**) Graphs show frequencies of live proliferating CFSE^low^ (**D**) CD4 T cells and (**E**) CD8 T cells in autologous DC-T cell co-culture from four individual donors. (**F-G**) DCs were treated with 3 μg/mL isotype control (black), anti-HLA-ABC (orange), anti-HLA-DR (green) or anti-CD86 (maroon) blocking antibodies for 1 hour, and then exposed to 2 μg/mL WT, H84T or D133G and co-cultured with CFSE labelled autologous CD4 or CD8 T cells (1:10 ratio DC:T cells) for 5 days. Bar graphs show frequency of live proliferating CFSE^low^ (**F**) CD4 T cells and (**G**) CD8 T cells from four individual donors with mean ± SEM. (**H-I**) DCs were exposed to nothing or 2 μg/mL WT, H84T or D133G and cultured for 24 hours, then the supernatants were collected and added to CFSE labelled autologous CD4 or CD8 T cells and cultured for 3 days. Bar graphs show frequency of live proliferating CFSE^low^ (**H**) CD4 T cells and (**I**) CD8 T cells from four individual donors with mean ± SEM. Friedman test with Dunn’s multiple comparisons test was used to assess statistically significant differences at * p<0.05 (** p<0.01).

### H84T treatment of DCs results in higher IAV-specific T cell expansion in response to replicating virus

To assess the effect of H84T on the induction of virus-induced T cell proliferation, we next co-cultured autologous DCs and T cells in the presence or absence of replicating IAV (**Figure 2A**). IAV induced both CD4 (**Figure 2B**) and CD8 (**Figure 2C**) T cell proliferation in the absence of BanLec. As expected, the D133G treated cells displayed similar T cell proliferation as in the condition without BanLec (**Figure 2B-C**). In contrast, WT induced high levels of CD4 (**Figure 2B**) and CD8 (**Figure 2C**) T cell proliferation regardless of whether IAV was present or not, in line with its mitogenic capacity. Interestingly, in the presence of H84T, both CD4 (**Figure 2B**) and CD8 (**Figure 2C**) T cell proliferation was consistently significantly higher than in the IAV exposed co-cultures without BanLec or with D133G (**Figure 2B-C**). To estimate the level of T cell proliferation in response to IAV not associated with the mitogenic effect of BanLec, we compared the fold change CD4 and CD8 T cell proliferation by normalizing (dividing) proliferation in the no virus conditions across groups (**Figure 2D-E**). We found that the fold change IAV-induced CD4 and CD8 T cell proliferation was actually lowest in the WT condition (**Figure 2D-E**), which implies that the robust T cell expansion in the presence of WT was not antigen-specific. Interestingly, in the presence of H84T, the fold change of IAV induced CD4 and CD8 T cell proliferation was significantly higher compared to no BanLec or D133G treated cells (**Figure 2D-E**), suggesting that H84T does not impair DC antigen-presentation but in fact enhances T cell proliferation in response to replicating IAV.

**Figure 2.**
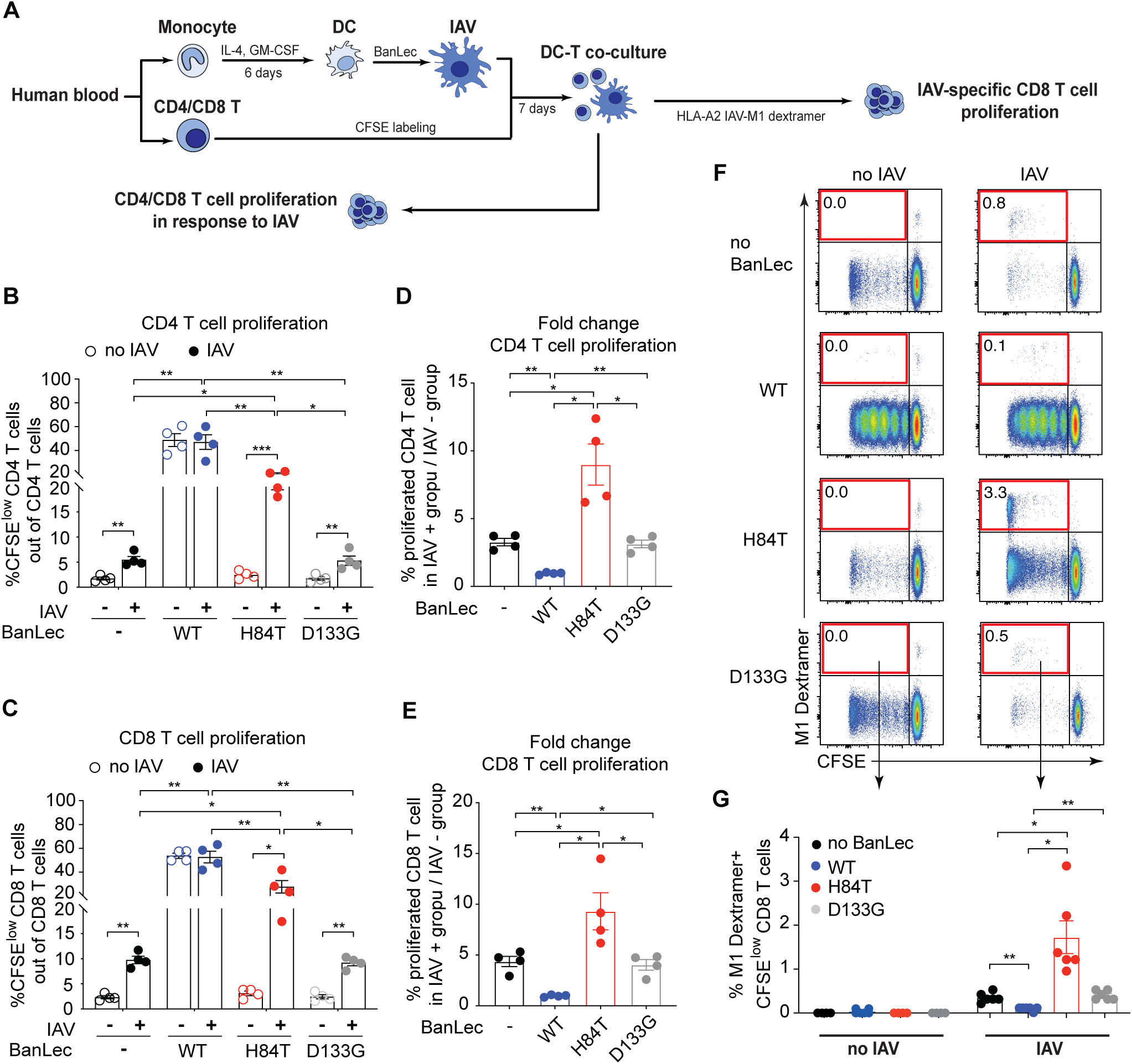
H84T treatment of DCs results in higher frequencies of IAV-specific CD8 T cells in response to replicating IAV. (**A**) DCs were exposed to nothing or 2 μg/mL WT, H84T and D133G for 1 hour and then exposed to no virus or 0.6 MOI IAV for 4 hours. Subsequently, CFSE labelled autologous CD4 or CD8 T cells were added (1:10 ratio MDDC:T cells) and co-cultured for 7 days. T cell proliferation was detected by CFSE dilution using flow cytometry. (**B-C**) Bar graphs show frequency of live proliferating CFSE^low^ (**B**) CD4 T cells and (**C**) CD8 T cells from four individual donors with mean ± SEM. (**D-E**) Bar graphs show the fold change of proliferated (**D**) CD4 T cells and (**E**) CD8 T cells with mean ± SEM, in which the frequencies of IAV induced CD4 and CD8 T cell proliferation were divided by no virus conditions. (**F-G**) DCs from HLA-A2 positive donors with detectable IAV-specific CD8 T cell memory responses were generated and exposed to nothing or to 2 μg/mL WT, H84T or D133G for 1 hour, and then exposed to no IAV or 0.6 MOI IAV for 4 hours. Subsequently, CFSE labelled autologous CD8 T cells were added (1:10 ratio DC:T cells) and co-cultured for 7 days. IAV M1-specific CD8 T cells were identified with an HLA-A2 Influenza A M1 dextramer and analysed using flow cytometry. (**F**) Dot plots show live CD8 T cells from one representative donor. The CFSE^low^ M1 Dextramer^+^ population (upper left quarter of the plots) represents proliferated IAV-specific CD8 T cells detected by IAV M1 Dextramer. Frequencies of CFSE^low^ M1 Dextramer^+^ CD8 T cells out of total CD8 T cells are displayed. (**G**) Bar graphs show frequency of live CFSE^low^ M1 Dextramer^+^ CD8 T cells from six individual donors with mean ± SEM. Statistical differences were assessed using RM one-way ANOVA with Tukey’s multiple comparisons test and considered significant at * p<0.05 (** p<0.01).

To study the effect of H84T on IAV-specific T cell responses, we generated DCs and isolated CD8 T cells from HLA-A2+ donors with detectable IAV M1-specific CD8 T memory cells to monitor proliferation of IAV-specific CD8 T cells in the presence of BanLec and/or IAV (**Figure 2A**). As expected, IAV M1-specific CD8 T cells proliferated (upper left quadrant) in response to IAV in cultures without BanLec or with D133G, while no IAV-specific CD8 T cell proliferation was observed in the absence of IAV (**Figure 2F-G**). In cultures with WT, no or little IAV M1-specific CD8 T cell expansion was observed in response to IAV (**Figure 2F-G**). Again, this supports the conclusion that the substantial T cell proliferation induced by the WT is mitogenic rather than virus-specific. Interestingly, we again observed that exposing DCs to H84T and IAV resulted in higher frequencies of proliferating IAV-specific CD8 T cells compared to co-cultures with no BanLec or D133G (**Figure 2F-G**). Together, this shows that the modified plant lectin H84T promotes superior virus-specific T cell responses to replicating IAV.

### H84T limits IAV infection in DCs, which preserves DC viability and allows expression of viral antigen for presentation to T cells

To understand why H84T treatment of IAV infected DCs results in generation of higher frequencies of virus-specific T cells, we studied DC-IAV-H84T interactions. We exposed DCs to replicating IAV in the presence or absence of BanLec and determined the viral infection by intracellular IAV nucleoprotein (NP) staining. The frequency of IAV NP+ DCs was significantly reduced in the presence of H84T and WT as compared to no BanLec or D133G treated groups (**Figure 3A-B**). Importantly, H84T also efficiently protected also human primary blood and tonsil myeloid DCs (mDCs) and plasmacytoid DCs (pDCs) from IAV infection (**Figure S1**).

**Figure 3.**
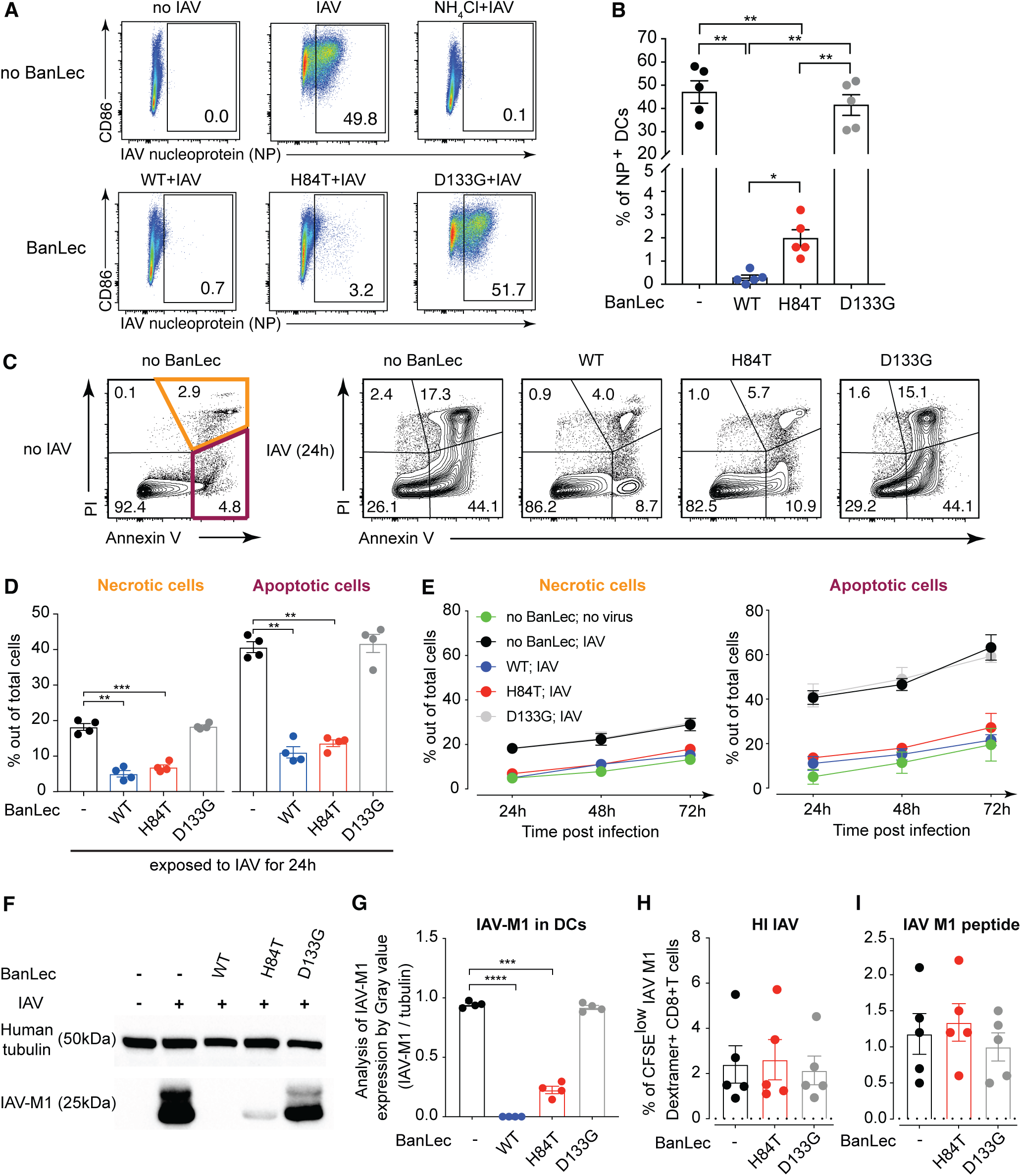
H84T effectively limits IAV infection in human DCs, and consequently preserves DC viability but allows expression of viral antigen for presentation to T cells. (**A-B**) DCs were exposed to nothing or 2 μg/mL WT, H84T or D133G for 1 hour, and then exposed to no IAV, or 0.6 MOI IAV with or without 20 mM NH_4_Cl for 24 hours to prevent viral fusion. (**A**) Dot plots show live DCs and numbers depict the frequency of IAV nucleoprotein (NP)+ DCs from one representative donor. (**B**) Graphs show frequency of IAV NP+ DCs from five individual donors with mean ± SEM. (**C-E**) DCs were exposed to nothing or 2 μg/mL WT, H84T or D133G for 1 hour, then exposed to no IAV or 0.6 MOI IAV for 24, 48 or 72 hours. Viability of DCs was detected by Annexin V and PI staining using flow cytometry. (**C**) Dot plots show DCs and numbers depict the frequency of PI or Annexin V positive DCs from one representative donor out of four. (**D**) Graphs show frequency of necrotic (orange) and apoptotic (maroon) cells out of total cells after 24 hours of exposure to IAV in four donors with mean ± SEM. Paired t test was used to analyze the data and data considered significant at * p<0.05 (** p<0.01, *** p<0.001). (**E**) Line graphs show the frequency of necrotic and apoptotic cells with mean ± SEM in four individuals. (**F-G**) IAV M1 protein in cell lysates from DCs exposed to nothing, 2ug/mL BanLec (WT, H84T or D133G) with or without 0.6 MOI IAV for 24 hours were determined by Western blot. Tubulin was used as the loading control. (**F**) One representative donor from four is shown. (**G**) Graph shows the analysis of IAV-M1 expression level by gray value in DCs exposed to replicating IAV. (**H-I**) DCs from HLA-A2 positive donors with detectable IAV-specific CD8 T cell memory responses were generated and exposed to nothing or to 2 μg/mL H84T or D133G BanLec for 1 hour, and then exposed to heat-inactivated IAV (HI IAV) or IAV M1 peptide (GILGFVFTL) for 4 hours. Subsequently, CFSE labelled autologous CD8 T cells were added and co-cultured for 7 days. IAV M1-specific CD8 T cells were identified with an HLA-A2 Influenza A M1 dextramer and analysed using flow cytometry. T cell proliferation was detected by CFSE dilution. Graph shows frequencies of live CFSE^low^ M1 Dextramer+ CD8 T cells with (**H**) HI-IAV and (**I**) IAV M1 peptide stimulation with mean ± SEM values from five individual donors. Dotted lines show the mean frequencies of live CFSE^low^ M1 Dextramer+ CD8 T cells in DC-CD8 T cell co-culture without HI IAV or IAV M1 peptide stimulation. Statistical differences were assessed using the Friedman test with the Dunn’s multiple comparisons test and considered significant at * p<0.05.

In response to pathogens including viruses, DCs undergo maturation characterized by upregulation of HLA-ABC (MHC I) and HLA-DR (MHC II), co-stimulatory molecules CD86/CD80 and CD40 to facilitate antigen-presentation and T cell interaction ^23 24^. Therefore, we next tested the impact of H84T on the MHC and co-stimulatory molecule expression of DCs during IAV infection. As expected, IAV itself was a potent inducer of DC maturation, while the different BanLec variants did not induce DC maturation on their own (**Figure S2A-D**). In line with their ability to inhibit IAV infection, the presence of WT or H84T resulted in less DC maturation as assessed by HLA-ABC, HLA-DR, CD86 and CD40 expression in response to IAV (**Figure S2A-D**). Similarly, replicating IAV induced secretion of TNFα and IL-6 from DCs but not in the presence of WT and H84T that inhibit infection (**Figure S2E-F**). Together, these findings show that greater DC maturation or higher cytokine secretion do not explain why H84T treatment results in superior propagation of virus-specific T cells.

We next determined the viability of IAV exposed DCs in the presence or absence of BanLec since IAV replication typically results in cytopathic effects, which in turn may affect the ability of DCs to interact with T cells. We used Annexin V and propidium iodide (PI) staining to identify apoptotic and necrotic cells respectively in the cultures after 24-72 hours of IAV exposure (**Figure 3C-E**). After 24 hours of exposure to IAV, in the presence of both WT and H84T the frequencies of apoptotic and necrotic DCs were significantly lower compared to no BanLec (**Figure 3C-D**). WT and H84T treatment preserved DC viability 48h and 72h post infection (**Figure 3E**), showing that the antiviral property of both H84T and WT coincides with preserved viability of DCs in the presence of replicating IAV. Still, only H84T treatment, but not WT, resulted in higher frequencies of IAV-specific T cells. Therefore, we next determined viral antigen load in DCs under different culture conditions since that is another rate limiting step in antigen presentation and T cell activation. Cell lysates from IAV exposed DCs displayed a strong IAV M1 protein band when cultured without BanLec or with D133G (**Figure 3F-G**). In lysates of DCs exposed to H84T and IAV, a weak but visible M1 protein band was detected, while no M1 protein was detected in lysates of WT treated DCs (**Figure 3F-G**). These data show that H84T treatment, but not WT, allows limited but detectable viral antigen expression in IAV exposed DCs.

Taken together, H84T significantly limits IAV infection in DCs, which preserves DC viability but allows some viral antigen to be translated and available for T cell presentation. We propose that this explains why H84T treatment of DCs in the presence of replicating IAV results in higher frequencies of IAV-specific T cells as compared to untreated DCs (more cell death) or DCs treated with WT (no viral antigen and non-specific T cell proliferation). To test this hypothesis, we added non-replicating forms of IAV antigen to the DC-T cell co-cultures: heat-inactivated IAV (HI IAV) that can bind and fuse but not replicate, or IAV M1 peptide (GILGFVFTL). HI IAV (**Figure 3H**) and IAV M1 peptide (**Figure 3I**) could both induce similar IAV M1-specific CD8 T cell proliferation in co-cultures regardless of treatment (**Figure 3H-I**). Taken together, we found that H84T treatment improves virus-specific T cell responses in the presence of replicating IAV and, importantly, does not impair immune responses to non-replicating IAV antigens.

### H84T treatment of IAV infected DCs rescues their capacity to initiate specific CD8 T cell responses against other antigens

IAV infection predisposes patients to secondary bacterial infection, which increases the risk of severe or fatal disease ^25,26^. We have previously reported that IAV infection of DCs impairs their ability to present antigen to CD8 T cells, which may contribute to increased susceptibility to secondary infections ^5^. To assess the impact of H84T treatment of IAV exposed DCs on mounting T cell responses against non-IAV antigens, we used cells from HLA-A2+ donors with detectable CD8 T cell memory responses against CMV pp65 protein, which was added as a second antigen in autologous DC and CD8 T cell co-cultures exposed to replicating IAV or HI IAV in the absence or presence of BanLec (**Figure 4A**). CMV pp65-specific CD8 T cells did not proliferate in the absence of CMV pp65 protein (**Figure S3**). As previously reported, DCs exposed to replicating IAV induced lower frequencies of CMV-specific CD8 T cells compared to DC not exposed to IAV or exposed to HI IAV (**Figure 4B-C**). As expected, D133G treatment generated similar results as did no BanLec (**Figure 4B** and **D**). Interestingly, H84T treatment resulted in significantly higher level of CMV-specific CD8 T cell proliferation by IAV infected DCs, as compared to IAV infected DCs exposed to D133G or not treated with BanLec (**Figure 4B** and **E**). In fact, in the presence of H84T, DCs exposed to replicating IAV induced similar CMV-specific CD8 T cell responses as did uninfected DCs or HI IAV exposed DCs (**Figure 4B** and **F**). Together, our data suggest that H84T rescues the impaired capacity of IAV infected DCs to induce CD8 T cell proliferation against other antigens, to comparable levels as seen in the absence of IAV infection.

**Figure 4.**
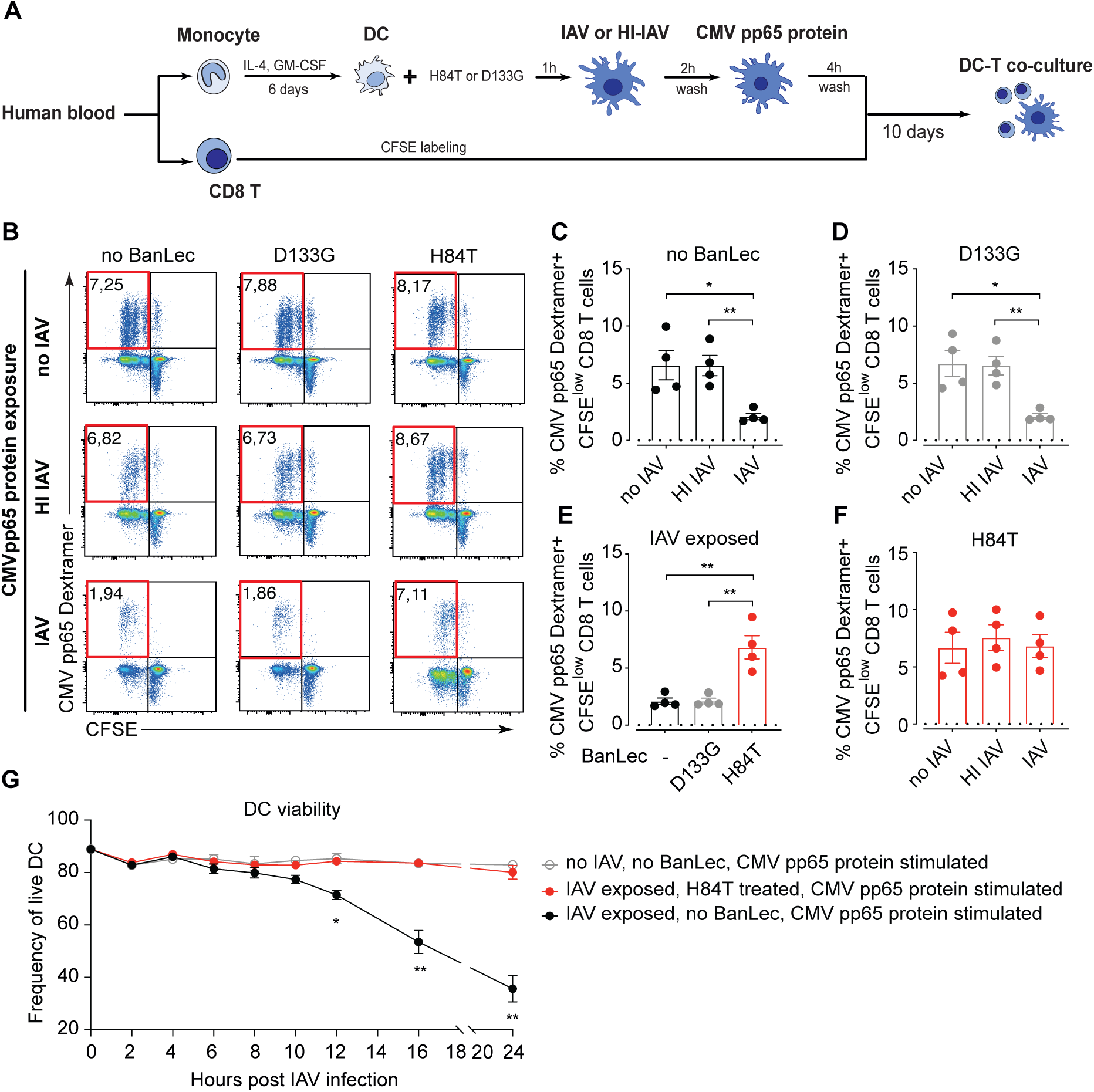
H84T restores the capacity of DCs to initiate pathogen-specific CD8 T cell expansion against CMV antigen post replicating IAV infection. (**A**) DCs differentiated from HLA-A2+ donors were pre-treated with nothing or H84T or D133G for 1h, and then exposed to IAV (0.6 MOI) or HI-IAV (0.6 MOI) for 2h. Then the cells were washed and the recombinant cytomegalovirus (CMV) pp65 protein were added as a second source of antigen. After 4h incubation, DCs were washed and co-cultured with autologous CFSE labelled CD8 T cells for 10 days. CMV-specific CD8 T cells were identified with an HLA-A2-CMV pp65-dextramer and analysed using flow cytometry. (**B**) Dot plots show live CD8 T cells from one representative donor. The CFSE^low^ CMV pp65 Dextramer^+^ population (upper left quarter of the plots) represents proliferated CMV-specific CD8 T cells. Frequencies of CFSE^low^ CMV pp65 Dextramer^+^ CD8 T cells out of total CD8 T cells are displayed. Bar graphs show frequency of live CFSE^low^ CMV pp65 Dextramer^+^ CD8 T cells in conditions (**C**) without BanLec, or in the presence of (**D**) D133G or (**F**) H84T from four individual donors with mean ± SEM. (**E**) Bar graphs show frequency of live CFSE^low^ CMV pp65 Dextramer^+^ CD8 T cells in DC-CD8 T cell co-culture with replicating IAV exposure from four individual donors with mean ± SEM. (**C-F**) Dotted lines show the mean frequency of live CFSE^low^ CMV pp65 Dextramer^+^ CD8 T cells in DC-CD8 T cell co-culture without CMV pp65 protein stimulation. (**G**) DCs differentiated from HLA-A2+ donors were pre-treated with nothing or H84T for 1h, and then exposed to nothing or IAV (0.6 MOI) for 2h. Then the cells were washed and the recombinant CMV pp65 protein added as a second source of antigen. The viability of the DCs was detected by Annexin V and PI staining using flow cytometry at 0, 2, 4, 6, 8, 10,12,16 and 24h post IAV infection. Line graph shows the frequency of live DCs out of total DCs from four individual donors with mean ± SEM. Statistical differences were assessed using RM one-way ANOVA with Tukey’s multiple comparisons test and considered significant at * p<0.05 (** p<0.01).

To understand the mechanism of action behind this observation, we determined the viability of DCs longitudinally after exposure to replicating IAV or no IAV, in the presence of CMV pp65 protein. We found that, without BanLec treatment, the viability of DCs exposed to replicating IAV started to significantly decrease 12 hours post infection as compared to DCs not exposed to IAV (**Figure 4G**). In the presence of H84T, the viability of DCs was comparable to that in the conditions with no IAV (**Figure 4G**). Thus, H84T likely rescues antigen-presenting capacity of DCs exposed to replicating IAV by preserving DC viability.

To sum up, our study strongly supports the development of H84T as novel antiviral against IAV. H84T not only directly blocks IAV replication, but further facilitates robust pathogen-specific CD8 T cell responses against not only influenza, but also against antigens from another pathogen after IAV infection. In addition, H84T holds promise to reduce the risk of severe disease by preventing the excess inflammatory cytokine production induced by influenza. In all, H84T treatment may enable patients to not only handle IAV but also better manage pathogens that can cause secondary infections that are the hallmarks of severe or even fatal influenza disease.

## DISCUSSION

Since its identification in 1990 ^27^, BanLec has been studied as a potential broad-spectrum antiviral due to its strong binding affinity for high mannose found on many enveloped viruses ^28^. However, systemic inflammatory responses in virus infection mouse models ^29^, driven by mitogenic activation and proliferation of T cells, has prevented the exploration of unmodified WT for clinical use. In contrast, the genetically engineered H84T is well tolerated in mouse models and does not induce T cell proliferation in human PBMCs ^17^. H84T has potent antiviral effect and protects mice from lethal influenza infection, which supports continued development for therapeutic applications ^12,17^. Still, the immunological mechanism of T cell proliferation, observed with WT but not with H84T, was unclear. Therefore, in this study, we first determined the impact of BanLec on human T cells. Interestingly, we found that WT did not induce proliferation by directly interacting with T cells. Instead, the mitogenic effect of WT required MHC II- and CD86-dependent APC-T cell interactions to induce T cell proliferation. This mechanism is similar to what has been described for superantigens such as staphylococcal enterotoxin A ^30^. While T cells did not proliferate in the presence of H84T, even when co-cultured with DCs that are professional APCs. Thus, our data confirm the absence of mitogenic effect of genetically engineered H84T on human T cells.

In response to viral infections like influenza, activation and expansion of virus-specific T cells are critical for effective viral clearance and long-lived immunological memory. This requires well-orchestrated cellular interactions between T cells and APCs, especially DCs. Upon IAV infection, DCs sense and capture the virus for subsequent antigen processing and presentation ^31–33^. However, IAV infection of DCs has been shown to impair their capacity to activate T cells ^5^. Thus, we determined the impact of H84T on the ability of DCs to support IAV-specific T cell expansion in response to replicating IAV. Interestingly, we found that H84T treatment resulted in more efficient expansion of IAV-specific CD8 T cells, despite the antiviral properties of H84T that limit viral replication and thus viral antigen availability. To investigate the mechanism of this observation, we determined DC maturation and cytokine production, necessary for T cell activation and expansion ^34 35^, in response to IAV infection during H84T treatment. We found that H84T did not induce superior DC maturation or cytokine production as compared to D133G or untreated cells. This indicates that H84T does not improve antigen-presentation of DCs directly. Instead, H84T might contribute to increased expansion of IAV-specific CD8 T cells via indirect factors.

IAV replication typically has cytopathic effects on infected cells and results in cell death in a dose- and time-dependent manner ^36–41^. We found that both WT and H84T effectively limited viral replication, which resulted in preserved DC viability upon IAV exposure/infection. Importantly, H84T still allowed some viral protein production, providing sufficient antigen for DCs to present to and activate virus-specific CD8 T cells while limiting the cytopathic effects of IAV. This could explain why H84T but not WT treatment resulted in higher frequencies of proliferating IAV-specific T cells since WT prevented translation of any viral protein that could be presented to T cells. In addition, H84T treatment resulted in higher frequencies of proliferating IAV-specific CD8 T cells only in response to replicating IAV but not in response to non-replicating forms of the same antigen such as heat-inactivated IAV or IAV M1 peptide. In conclusion this suggests that H84T treatment efficiently limits viral replication resulting in maintained DC viability yet allowing some viral antigen to be available for presentation to virus-specific CD8 T cells. Thus, rather than actively promoting IAV-specific T cell proliferation in response to IAV infection, H84T sustains the capacity of DCs to initiate virus-specific T cell responses otherwise impaired by the cytotoxic effects of replicating IAV.

IAV infection of human primary DCs impairs their ability to present other antigens to CD8 T cells ^5^. This may contribute to the increased susceptibility of influenza patients to secondary infections associated with more severe clinical outcomes ^25,26^. We found the H84T could also restore the capacity of DCs to initiate robust pathogen-specific CD8 T cell expansion against CMV antigen during IAV infection. Here we used another viral antigen as our second antigen; in future studies it would be interesting to establish methods to assess if bacterial antigens are also more potently presented after H84T treatment, as bacterial infections are the typical secondary infection in influenza patients. By enabling the handling other pathogen-derived antigens in the presence of replicating IAV infection, our data indicate the H84T treatment may reduce the risk of severe disease in influenza.

In response to IAV infection, DCs produce inflammatory cytokines such as TNFα and IL-6 to initiate and modulate immune responses. However, too excessive cytokine production may contribute to cytokine storm, resulting in more severe disease ^42^. Higher TNFα and IL-6 levels in influenza patients have been shown to correlate with severe disease or fatal outcome ^43^. It is noticeable that H84T significantly decreased TNFα and IL-6 production of DCs induced by IAV without limiting virus-specific T cell expansion. Thus, our data also indicate the benefit of H84T treatment to reduce the risk of cytokine storm and progression of severe influenza.

Developing an antiviral therapy that efficiently limits viral replication and cytopathic effects but allows the induction and expansion of virus-specific immune responses would be a valuable addition to existing influenza vaccines and therapies. In summary, our data support the future development of H84T as a therapeutic agent for use in humans to manage IAV infection, where its antiviral effects coupled with its ability to enhance virus-specific immune responses could be highly beneficial. In addition, H84T may also reduce the risk of severe disease by preventing excessive inflammatory cytokine production and enabling individuals to handle other pathogen-derived antigens after IAV infection. H84T also displays broad-spectrum antiviral effects in mouse and hamster models and cell lines on a wide range of enveloped viruses including influenza B virus (IBV), human immunodeficiency virus (HIV), severe acute respiratory syndrome coronavirus 2 (SARS-CoV-2), Middle East respiratory syndrome coronavirus (MERS), hepatitis C virus (HCV), Ebola virus and herpesviruses ^12,17,44–48^. Future studies of the impact of H84T on human immune cell responses against those viruses would further justify the application of H84T as a broad-spectrum antiviral for use in humans.

## MATERIALS AND METHODS

### BanLec

WT and genetically engineered H84T and D133G were prepared in E.coli and purified as previously described ^17^. Briefly, cleared E.coli lysates were added to Ni-NTA agarose that has been equilibrated with IMAC-25 buffer. Then BanLec proteins were eluted via column and dialyzed against PBS using Slide-alyzer dialysis cassettes.

### Human Subjects

This study was approved by the local Ethical Review Board at Karolinska Institutet and performed according to the Declaration of Helsinki. Human tonsils were obtained from adult patients undergoing routine tonsillectomies to treat obstructive sleep apnea. Informed consent was obtained from all donors. Buffy coats were obtained from healthy blood donors at the blood bank of Karolinska University Hospital.

### Isolation and culture of cells

MDDCs were differentiated and subsets of human primary DCs and T cells were isolated as previously described ^5,49^. In brief, PBMCs were harvested after Ficoll-Paque Plus (GE Healthcare) gradient separation of buffy coats from healthy blood donors. Blood monocytes and T cells were enriched and obtained by counterflow centrifugal elutriation using RosetteSep human monocyte and T cell enrichment cocktail (StemCell Technologies), respectively. To differentiate MDDCs, monocytes were cultured with 40 ng/ml GM-CSF and 40 ng/ml IL-4 (both from R&D Systems) in complete medium, R10 (RPMI 1640 with 10% FCS, 1% L-glutamine and 1% penicillin/streptomycin) at 0.5 × 10^6^ cells/ml. R10 was replaced on day 3 and immature MDDCs were harvested on day 6.

Blood and tonsil myeloid dendritic cells (mDCs) and plasmacytoid dendritic cells (pDCs) were obtained by using CD1c^+^ Dendritic Cell Isolation Kit (Miltenyi Biotec) and the Diamond Plasmacytoid Dendritic Cell Isolation Kit II (Miltenyi Biotec) respectively, as previously described ^49^. Briefly, tonsils were mechanically disrupted and filtered to get a single-cell suspension. Tonsil mononuclear cells (TMCs) were harvested after Ficoll-Paque Plus (GE Healthcare) gradient separation of single-cell suspension. Then, TMCs or PBMCs were labelled with a mixture of biotin-conjugated antibodies against lineage markers, Fc receptors, myeloid or plasmacytoid markers. The non-target cells were first depleted and then the target cells were further isolated by using anti-biotin Micro Beads and magnetic columns.

### IAV

Influenza A/X31 (derived from influenza A/Aichi/2/68; H3N2) was propagated in chicken eggs, purified and concentrated on sucrose gradients (Virapur). The 50% tissue culture-infective does for IAV was determined by infecting a light monolayer of MDCKs in the presence of trypsin and monitoring the cytopathic effect. Virus was replication incompetent after heat-inactivation at 56°C for 0.5 hour.

### DC blocking assay

DCs, labelled with 0.25 μM CFSE (ThermoFisher Scientific), were treated with isotype control (purified mouse IgG, Biolegend), purified anti-HLA-ABC, anti-HLA-DR or anti-CD86 (all from Biolegend) blocking antibodies for 1 hour, and then exposed to WT, H84T or D133G and then co-cultured with CFSE labelled autologous T cells for 72 hours. T cell proliferation was detected by CFSE dilution using flow cytometry (LSRFortessa, BD Biosciences).

### BanLec pre-treatment and IAV infection of DCs

DCs were pre-treated with nothing or WT, H84T or D133G for 1h, and then exposed to IAV for 24h. IAV infection was monitored using an anti-nucleoprotein (NP) antibody (Abcam) and flow cytometry. To prevent IAV infection, 20 mM NH_4_Cl was added to DCs before they were exposed to IAV.

### DC phenotype and cytokine secretion

After IAV infection, DCs were harvested, washed with PBS and surface stained with antibodies against CD1a, CD11c, CD14, HLA-DR, CD40, CD86, and CCR7 (all from BD Biosciences). Then DCs were washed, fixed, and analysed by flow cytometry. Supernatants were harvested and cytokines were measured by ELISA (TNFα and IL-6, R&D Systems).

### DC viability after IAV infection

DCs were pre-treated with nothing or WT, H84T and D133G for 1h, and then exposed to IAV. DCs were harvested after 24, 48 and 72h respectively, washed in ice-cold PBS twice and resuspended in 1X Annexin binding buffer, and stained with Annexin V and propidium iodide (PI) (ThermoFisher Scientific), and analysed by flow cytometry within 0.5h of processing.

### DC presentation of IAV to CD8 T

DCs differentiated from HLA-A2+ donors were pre-treated with nothing or WT, H84T or D133G for 1h, and then exposed to IAV (0.6 MOI), HI-IAV (0.6 MOI) or IAV M1 peptide (GILGFVFTL, 0.5 μg/mL) for 24h, and then co-cultured with CFSE labelled autologous CD8 T cells. After 7 days, cells were harvested and stained with HLA-A2 Influenza M1 (GILGFVFTL) dextramer (Immudex) for 15 min at room temperature followed by labelling with antibodies against CD3, CD4, CD8, CD69, CD14, CD19 and CD56 (all BD Biosciences), fixation and analysis by flow cytometry.

### DC presentation of CMV pp65 protein to CD8 T

DCs differentiated from HLA-A2+ donors were pre-treated with nothing or H84T or D133G for 1h, and then exposed to IAV (0.6 MOI) or HI-IAV (0.6 MOI) for 2h. Then the cells were washed and the recombinant cytomegalovirus (CMV) pp65 protein (ThermoFisher Scientific) was added as a second source of antigen. After a 4 h incubation, DCs were washed and co-cultured with autologous CFSE labelled CD8 T cells. After 10 days, cells were harvested and stained with HLA-A2-CMV pp65-dextramer (Immudex) for 15 min at room temperature followed by labelling with antibodies against CD3, CD4, CD8, CD69, CD14, CD19 and CD56 (all BD Biosciences), fixation and analysis by flow cytometry.

### IAV antigen load in DCs

After pre-treatment with nothing or WT, H84T or D133G for 1h, followed by IAV exposure for 24h, DC were harvested and lysed in SDS lysis buffer (Sigma-Aldrich). DNA was sheared mechanically and lysates were snap frozen on dry ice. Lysates were run on 4-12% Bis-Tris reducing gel, transferred to a PVDF membrane and blotted for viral proteins with the anti-IAV M1 monoclonal antibody (Abcam). Human tubulin was used as a loading control.

### Statistical analysis

Data were analysed using Prism 9 (GraphPad). Grouped data are generally presented as mean ± SEM. Comparisons between variables were performed using the Friedman test with Dunńs multiple comparisons test or paired t test where appropriate. A significance level of 95% was used and all tests were two-tailed.

## Author contributions

Experimental study design: M.Y., D.M.M. and A.S-S. BanLec generation: M.L. HLA-A2^+^ PBMCs screening: E.S. Data generation: M.Y., A.L., F.B. and S.L. Analysis and interpretation of data: M.Y. and A.S-S. Preparation of figures: M.Y. Manuscript writing: M.Y. and A.S-S. Critical revision of the manuscript: all authors.

## Acknowledgments

We thank the volunteers who have contributed to this study. We would like to thank Sang Liu for flowcytometry supervision. Work in the laboratory of D.M.M. was supported by the Forbes Institute of the Rogel Cancer Center at the University of Michigan, the University of Michigan MTRAC (Michigan Translational Research and Commercialization) organization, and R01AI175174 from the National Institutes of Health.

