## Supplemental Text and Figures for "A genetically engineered therapeutic lectin inhibits human influenza A virus infection and sustains robust virus-specific CD8 T cell expansion"

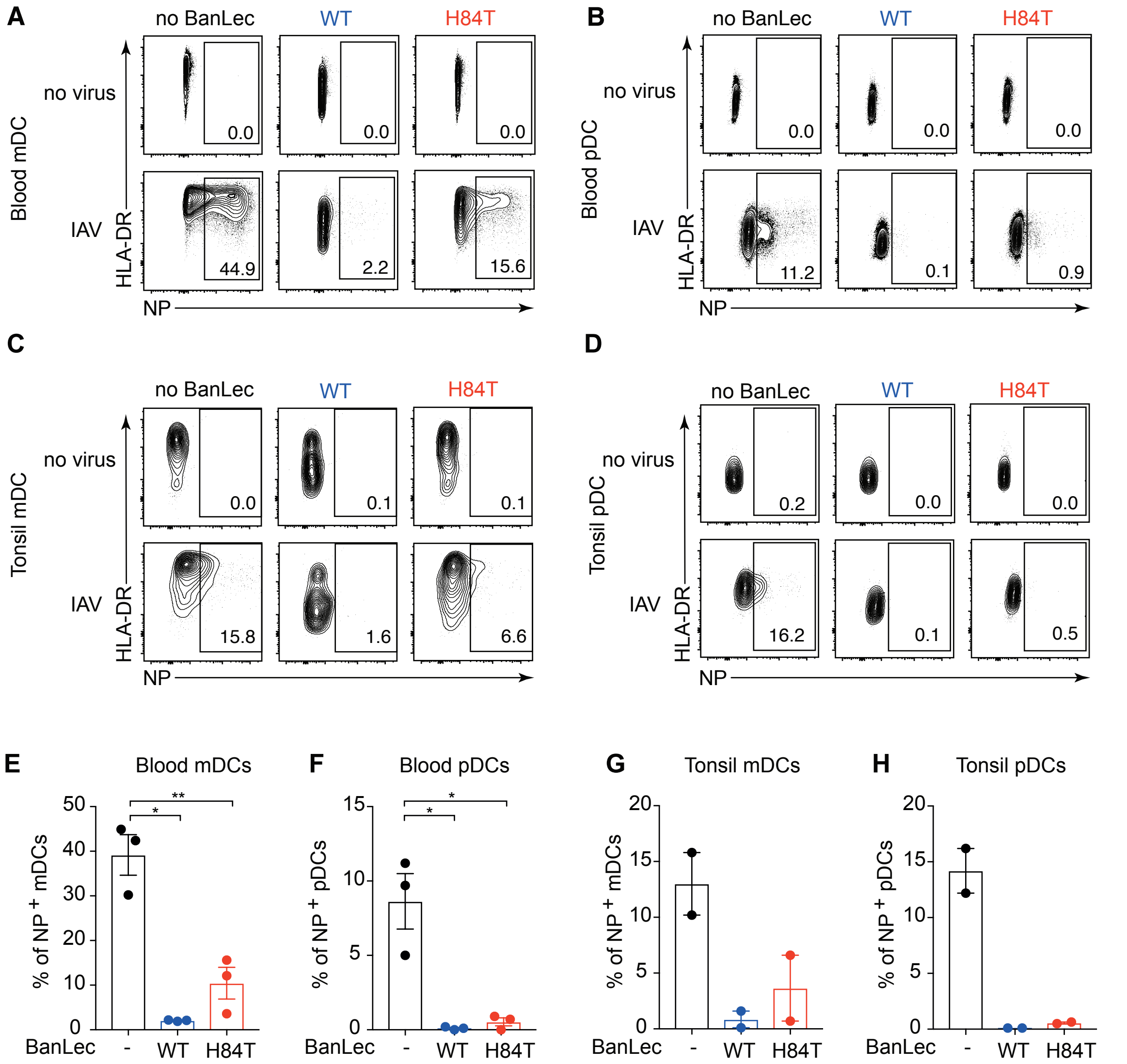


**Figure S1. WT and H84T inhibit IAV infection of human primary myeloid dendritic cells (mDCs) and plasmacytoid dendritic cells (pDCs) from both blood and tonsils.** pDCs and CD1c+ mDCs were isolated from human blood and tonsil. DCs were exposed to nothing or 2 µg/mL WT or H84T for 1 hour, and then exposed to no virus or 0.6 MOI IAV for 24 hours. (A) Dot plots show live blood mDCs and numbers depict the frequency of IAV NP+ mDCs from one representative donor out of three. (B) Dot plots show live blood pDCs and numbers depict the frequency of IAV NP + pDCs from one representative donor out of three. (C) Dot plots show live tonsil mDCs and numbers depict the frequency of IAV NP + mDCs from one representative donor out of two. (D) Dot plots show live tonsil pDCs and numbers depict the frequency of IAV NP + pDCs from one representative donor out of two. Graphs show frequency of IAV NP+ (E) blood mDCs, (F) blood pDCs, (G) tonsil mDCs and (H) tonsil pDCs with mean ± SEM. (E-F) Paired t test was used to assess statistically significant differences at * $p<0.05$ (** $p<0.01$).

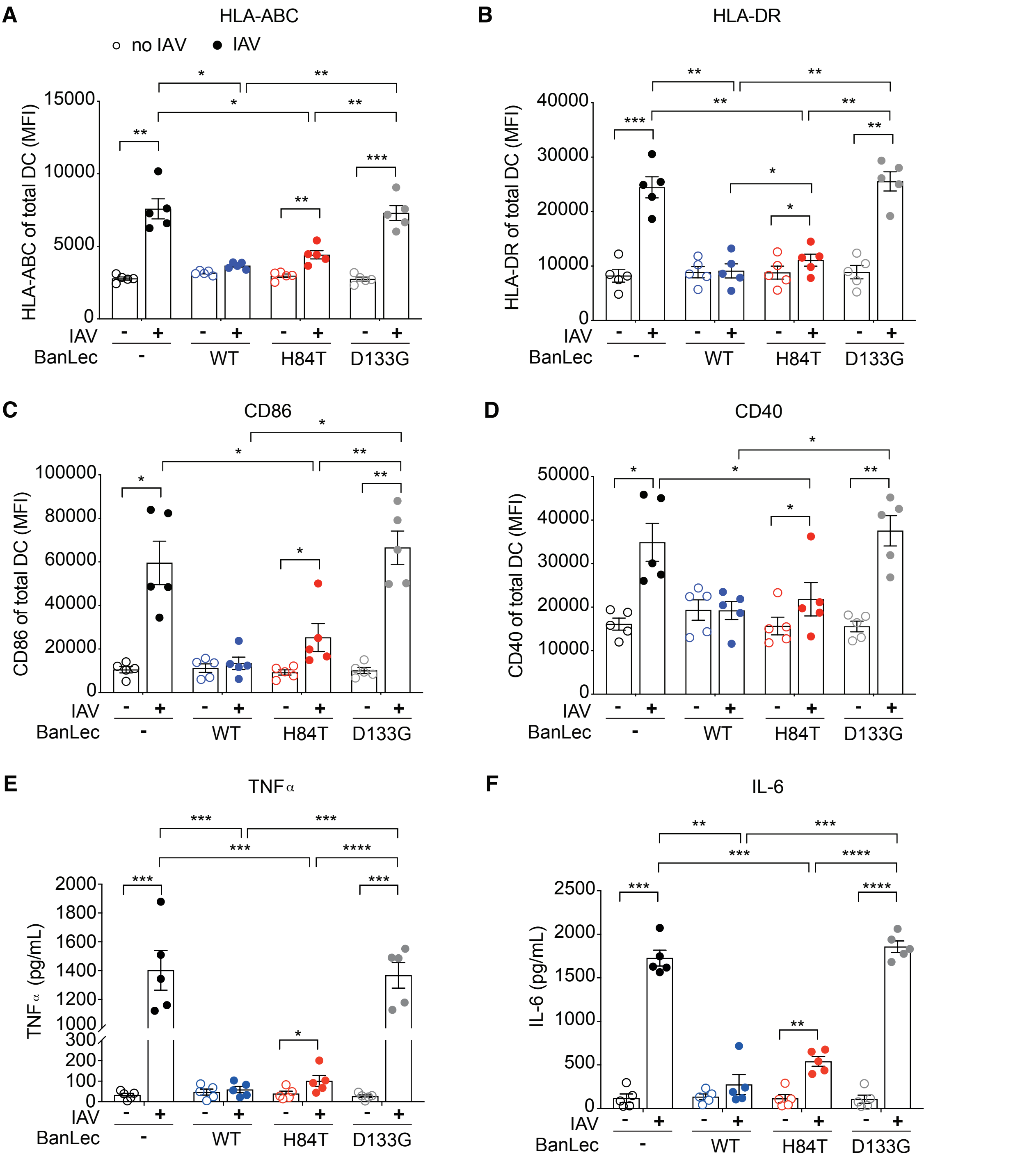


**Figure S2. WT and H84T prevent IAV-induced maturation of DCs.** DCs were exposed to nothing or 2 μg/mL WT, H84T or D133G for 1 hour and then exposed to no virus or 0.6 MOI IAV for 24 hours. The surface expression of (A) HLA-ABC, (B) HLA-DR, (C) CD86 and (D) CD40 on DCs in the absence or presence of different types of BanLec without IAV after 24 hours was measured by flow cytometry and displayed as geometric mean fluorescence intensity (MFI) from five individual donors with mean ± SEM. Supernatants were collected and the concentration of TNFα and IL-6 was determined by ELISA. Graphs show the concentration of (E) TNF and (F) IL-6 from five donors with mean ± SEM. RM one-way ANOVA with Tukey´s multiple comparisons test was used to assess statistically significant differences at * $p<0.05$, ** $p<0.01$, *** $p<0.001$, **** $p<0.0001$.

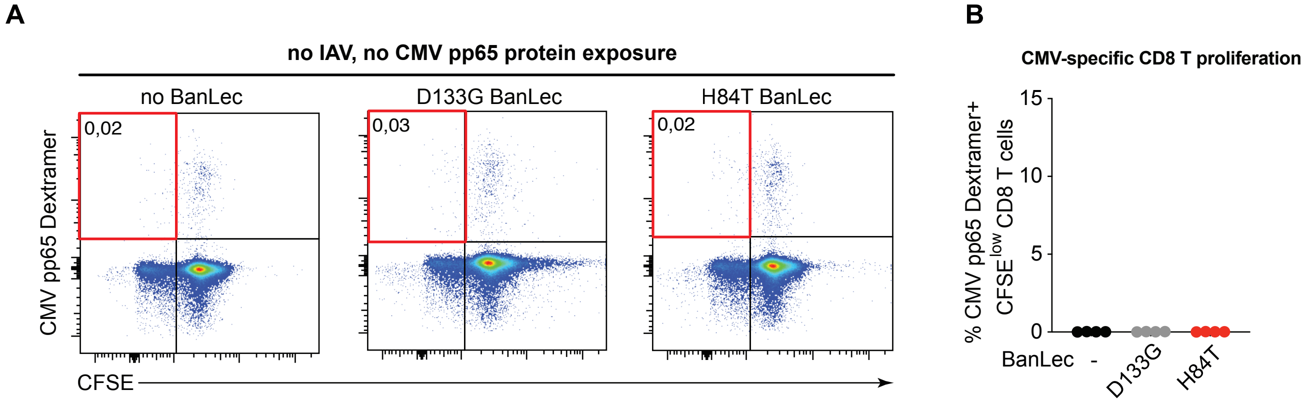


Figure S3. **Frequency of CMV-specific CD8 T cell proliferation in autologous DC co-cultures without CMV pp65 protein stimulation.** DCs differentiated from HLA-A2+ donors were pre-treated with nothing or H84T or D133G for 1h. Then the cells were washed without IAV or CMV pp65 exposure, and co-cultured with autologous CFSE labelled CD8 T cells for 10 days. CMV-specific CD8 T cells were identified with an HLA-A2-CMV pp65-dextramer and analysed using flow cytometry. (A) Dot plots show live CD8 T cells from one representative donor. The $CFSE^{low}$ CMV pp65 Dextramer+ population (upper left quarter of the plots) represents proliferated CMV-specific CD8 T cells. Frequencies of $CFSE^{low}$ CMV pp65 Dextramer+ CD8 T cells out of total CD8 T cells are displayed. (B) Bar graphs show frequency of live $CFSE^{low}$ CMV pp65 Dextramer+ CD8 T cells in conditions without BanLec, or in the presence of D133G or H84T from four individual donors with mean ± SEM. Statistical differences were assessed using RM one-way ANOVA with Tukey´s multiple comparisons test and considered significant at * $p<0.05$.
